# Detection of circular RNAs and their potential as biomarkers predictive of drug response

**DOI:** 10.1101/2023.01.08.522775

**Authors:** Julia Nguyen, Anthony Mammoliti, Sisira Kadambat Nair, Emily So, Farnoosh Abbas-Aghababazadeh, Christoper Eeles, Ian Smith, Petr Smirnov, Housheng Hansen He, Ming-Sound Tsao, Benjamin Haibe-Kains

## Abstract

The introduction of high-throughput sequencing technologies has allowed for comprehensive RNA species detection, both coding and non-coding, which opened new avenues for the discovery of predictive and prognostic biomarkers. However the consistency of the detection of different RNA species depends on the RNA selection protocol used for RNA-sequencing. While preliminary reports indicated that non-coding RNAs, in particular circular RNAs, constitute a rich source of biomarkers predictive of drug response, the reproducibility of this novel class of biomarkers has not been rigorously investigated. To address this issue, we assessed the inter-lab consistency of circular RNA expression in cell lines profiled in large pharmacogenomic datasets. We found that circular RNA expression quantified from rRNA-depleted RNA-seq data is stable and yields robust prognostic markers in cancer. On the other hand, quantification of the expression of circular RNA from poly(A)-selected RNA-seq data yields highly inconsistent results, calling into question results from previous studies reporting their potential as predictive biomarkers in cancer. We have also identified median expression of transcripts and transcript length as potential factors influencing the consistency of RNA detection. Our study provides a framework to quantitatively assess the stability of coding and non-coding RNA expression through the analysis of biological replicates within and across independent studies.

## INTRODUCTION

Next-generation genomic sequencing technologies and advancements in computational methods have made it possible to deeply profile the transcriptome of cancer cells (1). More specifically, RNA-sequencing (RNA-seq) has allowed for profiling of coding RNA (mRNA), and non-coding RNA (ncRNA), which expand into a vast array of RNA species such as long non-coding RNA (lncRNA), small nucleolar RNA (snoRNA), and circular RNA (circRNA) (2, 3). However, RNA detection is impacted by RNA-seq selection protocols used to isolate the species of interest from ribosomal RNA (rRNA), which accounts for the bulk of cellular RNA (2). These protocols include isolating RNA with polyadenylated tails, such as protein-coding mRNA, and rRNA-depletion, which aims to remove rRNA from cellular RNA. Ribosomal RNA depletion is suggested to be better suited for isolating and detecting circRNAs as they do not possess poly-A tails, even though they may contain sequences enriched in adenine and therefore be detected in low quantities with poly(A)-selection (2, 4). Moreover, rRNA-depletion is often conjoined with RNAse-R to enrich for circRNAs in a sample before RNA-seq by cleaving linear RNA, along with RT-qPCR for validating RNA-seq detection (expression) through the use of specific primers for quantification (4–6).

Circular RNAs have been investigated for their potential role in carcinogenesis (7) and response/resistance to anticancer therapies in multiple cancer types (8–10). Their unique closed-loop structure allows them to resist degradation from RNA enzymes, making them more stable than their linear counterparts (11). Recent studies have used poly(A)-selected RNA-seq data for circRNA detection (12), and identified subsequent drug associations from pharmacogenomic datasets for biomarker discovery (7). However, the use of unsuitable circRNA isolation protocols (i.e., poly(A)-selection) (2, 4), the use of various publicly available detection tools, such as CIRI2 (13) and CIRCexplorer2 (4, 13), may result in divergent performance for circRNA quantification.

Moreover, with the discovery of many potential transcriptomic biomarkers across large pharmacogenomic studies (14), investigating the consistency of expression quantification of transcripts across inter-lab replicates is important for validating the discovery of these candidate biomarkers (15–17). Consequently, whether or not the predictive value of circRNAs for therapy response generalizes in independent datasets remains an open question, consequently hindering their translation into clinical assays.

To further investigate this issue, we examined the circular RNA space from the three largest pharmacogenomic datasets available to date using CIRI2 and CIRCexplorer2 for circRNA detection from poly(A)-selected RNA-seq data. We assessed the consistency of the quantification of circRNA, mRNA, and corresponding gene expression in 48 cancer cell lines across inter-lab replicates. Comparison of RNA-seq selection protocols was done with cell lines and validated on

51 lung adenocarcinoma patients with matched rRNA-depleted profiles. This approach enabled us to compare RNA-seq selection protocols for circRNA detection and for the detection of prognostic biomarkers. We used supervised learning to identify the genomic features that influenced mRNA transcript stability across inter-lab biological replicates. The relationship between the stability and the predictive value for drug response of these mRNA transcripts were also investigated. Our results indicate that the consistency of RNA-seq expression for circRNA detection is unreliable in poly(A)-selection when compared to rRNA-depletion, raising doubts about their reported predictive value for drug response *in vitro*.

## METHODS

### Pharmacogenomic datasets

We used the RNA-seq and pharmacological data from the largest preclinical pharmacogenomics studies, namely datasets from the Genentech Cell line Screening Initiative (gCSI) (18, 19), the Cancer Cell Line Encyclopedia (CCLE) (20–22), and the Genomics of Drug Sensitivity in Cancer (GDSC2) (23–25), as processed in the ORCESTRA platform (26). These datasets collectively include data for 1,602 unique cell lines, 205 unique drugs, and 233,921 drug sensitivity experiments (Supplementary Table 1) (26–29). In addition, these datasets include RNA-seq profiles for different sets of cancer cell lines, which all employ a poly(A)-selection protocol for RNA-seq with varying average number of reads. Importantly, these studies included the same 48 cell lines (from 12 tissue types), enabling the assessment of the consistency of expression quantification for circRNA, mRNA, and the corresponding genes.

### RNA detection

To address the stability of circRNA expression quantification, we assessed the consistency of inter-lab circRNA detection and compared it against overall gene and transcript (mRNA) expression as benchmarks (Supplementary Figure 1). We utilized Kallisto (v.0.46.1) for computing overall gene and transcript-specific abundance using Gencode v33 transcriptome, and CIRI2 (v.2.0.6) and CIRCexplorer2 (v.2.3.8) using default parameters for exploring circRNA detection between experimental protocols. Transcript log_2_+1 normalized counts were used. To obtain transcript and gene counts from Kallisto, we isolated protein-coding transcripts and filtered out ones that were: (*i*) housekeeping as determined by the Housekeeping and Reference Transcript Atlas (HRT Atlas) (30); (*ii*) were not expressed across all datasets and possessed a median expression of zero across all datasets, to enable a robust estimate of stability across replicates (see Stability Index in METHODS). A total of 19,754 genes and 78,643 isoform transcripts were included. circRNA transcripts, obtained from both CIRI2 and CIRCexplorer2, were filtered to keep only: *(i)* exonic circRNAs, and *(ii)* circRNA transcripts with at least 2 read counts. circRNA transcripts were then filtered to remove those not present in at least 12 of the 48 biological replicates, resulting in a total of 19 transcripts from CIRI2 and 21 transcripts from CIRCexplorer2. Nonlinear dimensionality reduction was performed using Uniform Manifold Approximation and Projection (UMAP) to visualize similarities in transcript quantification in cell lines across datasets.

### Consistency of expression quantification

Many transcripts have been identified as potential biomarkers predictive of drug response in large cell-line pharmacogenomic studies (14, 31). However, the consistency of transcript abundance estimates across inter-lab biological replicates remains an open question, even with the utilization of RNA-seq data with the same selection protocol, such as poly(A)-selection (15, 16). The potential lack of consistency may impact the biomarkers discovered, as the expression of a transcript and its associated drug response in one dataset could yield a completely different result from the same transcript-drug pair in another dataset. Therefore, addressing transcript abundance stability across replicates will provide increased confidence in the biomarkers discovered. The RNA stability (stability index; SI) between datasets was estimated using the Spearman correlation coefficient:

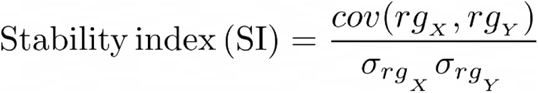

where cov(rgX, rgY) is the covariance of the rank variables (RNA log_2_+1 normalized transcript counts for two datasets), while σ_rgX_ σr_gY_ are the rank variable standard deviations. SI was computed across 48 biological replicates from gCSI, CCLE, and GDSC2 using log_2_+1 normalized transcript counts between dataset pairs (gCSI/CCLE, gCSI/GDSC2, GDSC2/CCLE). Transcripts with SI > 0.8 were considered to yield stable expression across replicates for the purpose of defining the binary classification.

### RNA-seq selection protocol

The detection of RNA species through RNA-seq is affected by the selection protocol used before sequencing (2), which has downstream consequences on biomarker discovery. RNAs with a polyadenylated (poly-A) tail represent 1-5% of eukaryotic cellular RNA and mostly consist of mRNA and lncRNA (32). They can be isolated from rRNA, which accounts for >80% of cellular RNA (2, 33), through enrichment by poly(A)-selection, which involves the binding of poly-A tails to oligo-dT molecules that are hybridized to beads (32). However, RNA without poly-A tails, such as circRNA, can only be detected in low quantities using poly(A)-selection, thus requiring a different isolation protocol from rRNA (4). A more suitable isolation method for circRNA is rRNA-depletion where only rRNA is eliminated from the rest of cellular RNA (32), a method that is employed by two technologies known as RiboZero and RiboMinus (34). Other methods for rRNA-depletion include the use of kits such as SMARTer Stranded Total RNA-Seq Kit v2 (Pico Input Mammalian), which are optimal for isolating low quantity RNA by preventing loss through removing rRNA cDNA generated after reverse transcriptase (35).

We compared two poly(A)-selected GDSC2 replicates (HCT 116; HeLa) with RiboZero profiles obtained from the Sequence Read Archive (SRA) (36) (Supplementary Table 2). We then compared three poly(A)-selected cell line circRNA profiles (22Rv1, LNCaP, and PC3) with matched RiboMinus profiles before and after RNAse-R treatment (37). Only circRNAs that were enriched by at least 1.5-fold in the matched RNAse R sample were retained. Lastly, poly(A)- selected and rRNA-depleted (SMARTer Stranded Total RNA-Seq Kit v2) RNA-seq data from 51 patient lung adenocarcinoma tumors were utilized for circRNA detection and experimental protocol comparison.

### Training sets for supervised learning

The use of supervised learning for identifying features that influence the prediction of mRNA consistency across inter-lab biological replicates holds the potential to provide insight into the biological and sequencing factors that impact the quantification of mRNA abundance. We therefore trained a linear regression model using a training set of labeled stable/unstable mRNA transcripts to identify transcript features with the greatest influence on mRNA stability. In order to determine the optimal number of mRNAs to include in the training set for supervised learning, the intersection of top stable/unstable mRNAs between the three dataset pairs across multiple quantiles (5% – 35%) was investigated. We used the Kuncheva Index (IC) to measure the significance of the intersections of stable and unstable mRNAs across the three datasets in a manner that also corrects for chance:

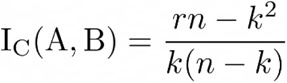

where A and B are feature subsets, r = |A ∩ B| (intersection of feature subsets), of the same size, k, while n is the total number of features (38). An IC of -1 indicates no similarity between feature sets, 0 indicates similarity of random chance, and 1 indicates identical feature sets. The quantile maximizing the Kuncheva index is selected to define the binary classification task for supervised learning.

### Feature space selection

mRNA expression and sequencing features were investigated, using supervised learning, for potential influence on mRNA stability across biological replicates. The genomic features included: (*i*) median expression; (*ii*) length (sum of length of exons); (*iii*) number of exons; (*iv*) GC-content; (*v*) 3’/5’ mapping bias; (*vi*) respective gene mappability. Median expression data was obtained by computing the median of log_2_+1 normalized counts for each mRNA across the 48 biological replicates. For each dataset pair, transcript median expression values from the remaining dataset was used (e.g., GDSC2 transcript median expression used for gCSI/CCLE pair). mRNA length and exon numbers were obtained through the *GenomicFeatures* Bioconductor R package (v.1.41.3), while GC-content was obtained through the *ensembldb* Bioconductor R package (v.2.13.1). 3’ mapping bias is often exhibited with poly(A)-selected data, as the poly-A tail at the 3’-end is selected for, including for degraded RNA (39), enriching for sequences at the 3’ ends.

5’ mapping bias can occur with oligo-DT primer use during cDNA generation, which can bind to internal adenine stretches, resulting in truncated 5’ cDNA (40). 3’/5’ bias was computed by using a formula derived from Picard’s CollectRnaSeqMetrics (broadinstitute.github.io/picard/), which is:

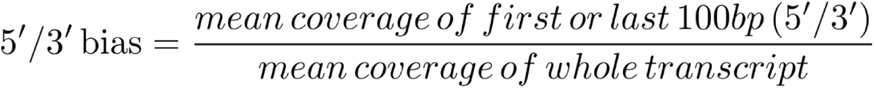

Coverage for the 3’/5’ bias calculations were computed using *mosdepth* (v.0.3.1). Lastly, respective gene mappability (single and multi-read) was computed using Umap (bismap.hoffmanlab.org/) (41).

### Estimating association between genomic features and stability

We trained a linear regression model using *caret* (v.6.0-92), where the training dataset consists of the mRNA expression and sequencing features, along with the SI as the dependent variable.

Mean squared error (MSE) was the performance scoring metric used for the linear regression models. MSE is the mean squared difference between actual and predicted stability values:’

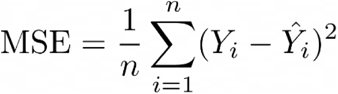

where 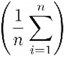 is the sum of a sequence of squares of error of *Y_i_* (observed values) - *Ŷ_i_* (predicted values), divided by *n*.

Ten-fold cross validation (CV) was performed with 10 repetitions, where the final performance is the mean ± standard deviation across all 10 folds and 10 repetitions. Each feature was then permuted (n=20,000) and another 10-fold CV was performed with 10 repetitions following each permutation. A scoring metric (MSE) was obtained from the 10-fold CV, where the mean score across all folds and repetitions was obtained for each permuted instance. The influence of each feature on predicting transcript stability (SI) was determined by taking the difference between the reported MSE scores from before and after the feature permutation (baseline_MSE_ - permuted_MSE_) (Supplementary Figure 2). This was repeated for each of the eight genomic features across all three dataset pairs.

### Transcriptomic stability and predictive value for drug response

In order to investigate the relationship between stability and predictive value for drug response, the area above the drug dose-response curve (AAC) was used as a drug response metric in order to measure cell viability across a range of drug concentrations (42). The CellTitre Glo assay was used in all three datasets to measure the cell viability in the control and drug experiments (18, 20, 23, 43). Predictive value for drug response was quantified using the concordance index (CI), computed for each gene/mRNA-drug pair as a meta-estimate across gCSI, CCLE, and GDSC2 using *survcomp* (v.1.40.0) Bioconductor R package. In addition, predictive value was also quantified with the Pearson correlation coefficient (18, 20, 23) and plotted as a forest plot with a 95% confidence interval using *forestplot* (v.1.1.0) R package. The transcripts were ranked by the mean SI across the dataset pairs, where only transcripts filtered for our supervised model were used. We assessed the relationship between transcript stability (randomized and non-randomized SI) and drug response predictive value using the rank-based Spearman correlation for each transcript used in our supervised model across all genes, where genes with less than 5 transcripts were discarded. Only drugs that intersected across the three datasets were selected.

### Survival analysis

Having demonstrated the best protocol for robust circRNA detection, we used this data to look for prognostic markers among the circRNA space available. Overall survival data was obtained for 51 patients with lung adenocarcinoma, where Kaplan- Meier plots were generated with the *survival* (v.3.2-7) and *survminer* (v.0.4.8) R packages, along with the *survcomp* (v.1.40.0) Bioconductor R package. In order to prevent software bias, only circRNAs detected by both CIRI2 and CIRCexplorer2 were retained, along with possessing a CI > 0.65 or < 0.35, and FDR < 0.05. Resulting circRNAs were validated with circBase (44).

### Research Reproducibility

To achieve full transparency and reproducibility, we shared all the curated data, computer code and software environment for readers to easily scrutinize, modify and reuse our research outputs. The pharmacogenomic datasets used are available on ORCESTRA (orcestra.ca) (26). The Zenodo digital object identifiers (DOI) for each data object are: gCSI (10.5281/zenodo.4737437); CCLE (10.5281/zenodo.3905461); GDSC2 (10.5281/zenodo.5787145). The computer code and snakemake pipelines for circRNA expression quantification via CIRI2 and CIRCexplorer2 are available on GitHub (https://github.com/bhklab/circRNA-detection). All components of data analysis, including the code and the data files, can be found in our Code Ocean capsule where the results can be reproduced directly from the platform (<u>codeocean.com/capsule/1660304/tree</u>).

## RESULTS

### Inconsistent circRNA quantification across datasets

The consistency of circRNA detection was investigated, along with the detection of mRNA and corresponding genes as benchmarks. Kallisto, CIRI2 and CIRCexplorer2 were used to quantify isoform/overall gene expression and circRNA expression, respectively, from poly(A)-selected RNA-seq data from 48 cell lines across three pharmacogenomic datasets (18, 25, 45). We first use UMAP (46) to visualize, in a 2-dimensional space, the local similarity between cell lines’ molecular profiles in the three pharmacogenomic datasets (Figure 1). We observed that, for isoform and overall gene expression, the biological replicates of cell lines were clustered closely, while cell lines were not clustered by studies. On the contrary, cell lines are strongly clustered by study rather than biological replicates when the circRNA expressions are used to compute similarities (Figure 1, Supplementary Figure 3a-d). The lack of similarity in the quantification of circRNA from cell lines across datasets suggests that circRNA detection is highly inconsistent across inter-lab replicates, supporting the concerns regarding the validity of circRNA studies highlighting their potential role as biomarkers for drug response.

**Figure 1.**
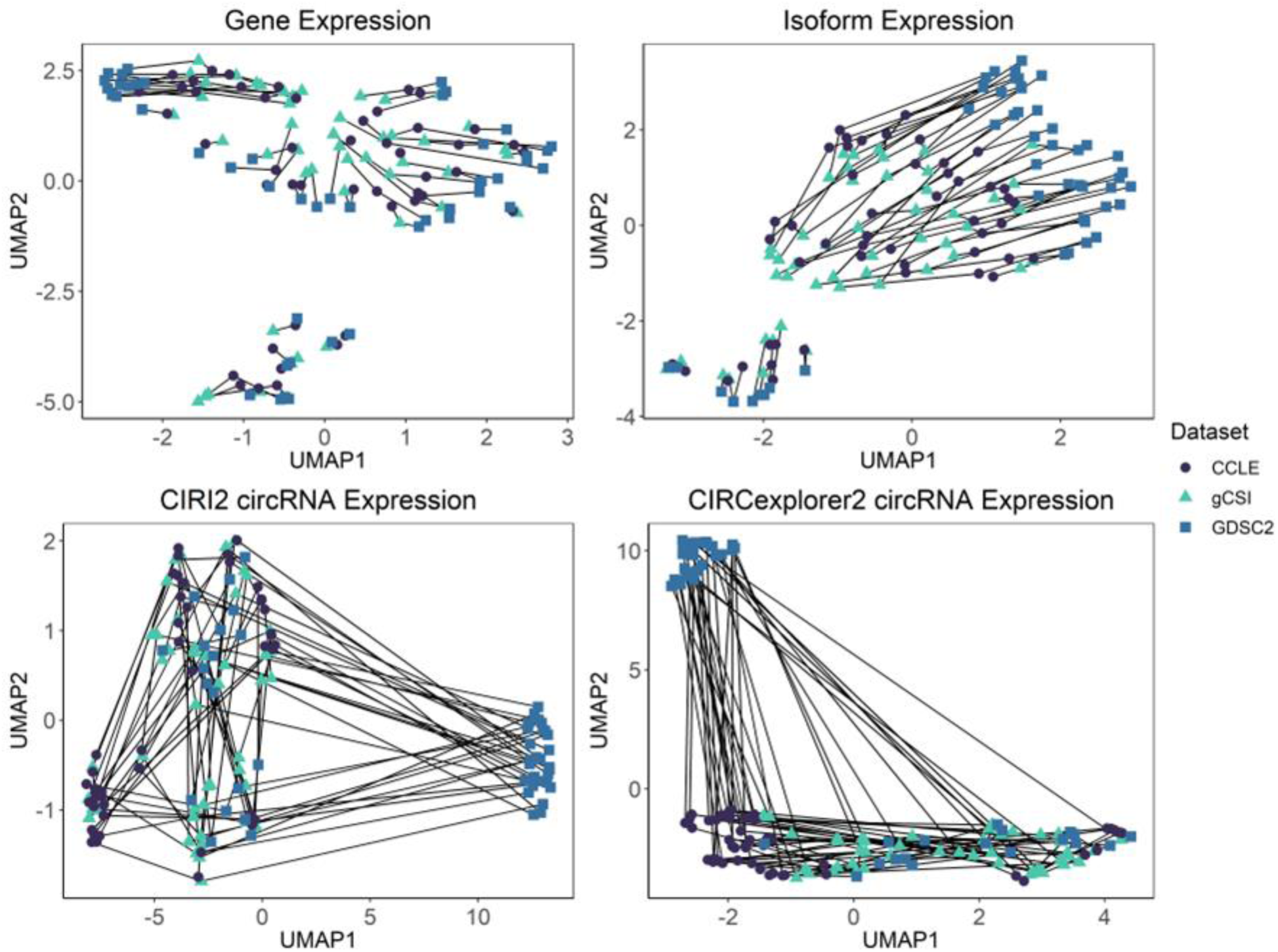
Transcript expression data dimensionality reduction by UMAP. Transcript expression across 48 biological replicates from 3 datasets (gCSI, CCLE, GDSC2). Expression is log_2_+1 normalized read counts. Lines connect individual cell lines from 3 datasets. Data points coloured and shaped by dataset (purple circles, CCLE; green triangles, gCSI; blue squares, GDSC2).

### Inconsistent transcript expression across datasets

Investigating the stability of RNA transcript abundance provides a further criteria for the evaluation of their potential as reliable biomarkers. The stability of circRNA transcripts, mRNA transcripts, and genes corresponding to the protein-coding transcripts were calculated across the dataset pairs – gCSI/CCLE, gCSI/GDSC, gCSI/CCLE. 283 circRNA transcripts were obtained from CIRI2 and 280 circRNA transcripts from CIRCexplorer2. From Kallisto, a total of 60,248 protein-coding transcripts were obtained. 19,957 protein-coding genes that matched with the previously obtained mRNA transcripts were used.

circRNA transcript expressions from both CIRI2 and CIRCexplorer2 were found to be highly unstable, with only 0.94% of CIRI2 transcripts and 1.4% of CIRCexplorer2 transcripts being classified as stable (SI > 0.8). Although the distribution of stability indices of mRNA transcripts were found to be better than random chance (p-value < 2.2e-16), only ∼5-8% of transcripts were classified as stable across replicates for each dataset pair, yielding highly unbalanced datasets. The distribution of the genes that correspond to the filtered mRNA transcripts was also found to be better than random chance (p-value < 2.2e-16) (Supplementary Figure 4), and ∼24% of the genes considered to be stable (Table 1).

**Table 1.**
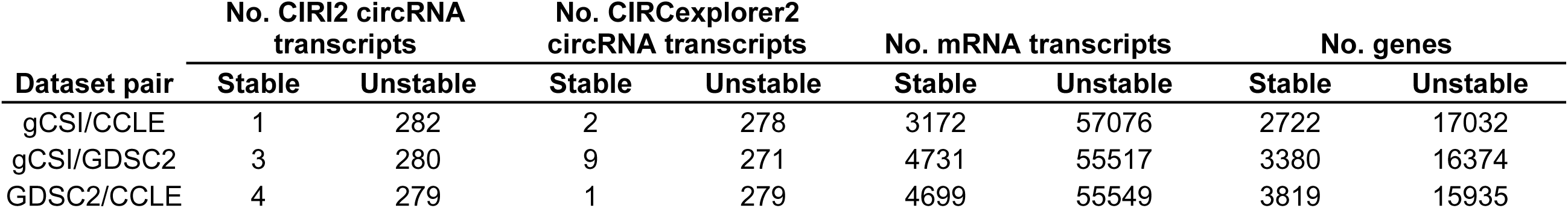
Number of stable and unstable transcripts expressed across biological replicates per dataset pair. Stability computed using the stability index. Stable transcripts possess a stability index > 0.8, while unstable transcripts possess a stability index < 0.8.

| Dataset pair | No. CIRI2 circRNA transcripts |  | No. CIRCexplorer2 circRNA transcripts |  | No. mRNA transcripts |  | No. genes |  |
| --- | --- | --- | --- | --- | --- | --- | --- | --- |
|  | Stable | Unstable | Stable | Unstable | Stable | Unstable | Stable | Unstable |
| gCSI/CCLE | 1 | 282 | 2 | 278 | 3172 | 57076 | 2722 | 17032 |
| gCSI/GDSC2 | 3 | 280 | 9 | 271 | 4731 | 55517 | 3380 | 16374 |
| GDSC2/CCLE | 4 | 279 | 1 | 279 | 4699 | 55549 | 3819 | 15935 |

The high percentage of unstable circRNA transcripts suggest they cannot be robustly quantified, and therefore are generally not suitable for biomarker discovery. These results also suggest that the stability of individual transcripts should be assessed before investigating their potential as biomarkers.

### RNA-seq with poly(A)-selection yields low circRNA detection

As the detection of circRNAs is affected by the RNA-seq selection protocol used (4), the consistency and accuracy of their profiling is at risk, which ultimately impacts biomarker discovery. To investigate the difference between poly(A)-selection and rRNA-depletion methods, two RiboZero and three RiboMinus cell lines were matched and compared to the corresponding poly(A)-selected cell lines. The RiboZero HCT 116 and HeLa cell lines yielded a 0.3 – 0.74-fold increase in the log_2_+1 normalized number of circRNA transcripts detected in comparison to their poly(A)-selected counterparts, however both CIRI2 and CIRCexplorer2 possessed similar detection profiles, with a 0.065-fold increase in circRNA counts in CIRI2 compared to CIRCexplorer2 (Supplementary Figure 5). When comparing the three poly(A)-selected cell lines (LNCaP, 22Rv1, PC3) with matched RiboMinus samples, there was a 4.9 – 16.7-fold increase in counts per million normalized circRNA count detected in the RiboMinus samples in samples without RNase R, and a 137 – 309-fold increase in samples with RNase R (Supplementary Figure 6). However, there was a 95% decrease in circRNA detection in the poly(A)-selected cell lines after validating them with their matched RNAse R samples(4). After validation through RNAse R, there was an 18.9 – 37.9% decrease in circRNA counts compared to isolation without RNAse R. To further investigate the suitability of poly(A)-selection protocol on circRNA detection, CIRI2 and CIRCexplorer2 were utilized to compare the circRNA profiles for 51 lung adenocarcinoma patient tumors with both poly(A)-selected and matched rRNA-depleted samples. Both CIRI2 and CIRCexplorer2 resulted in similar circRNA detection profiles, with a 1.3-fold increase in circRNA counts in CIRI2 compared to CIRCexplorer2. The rRNA depleted samples exhibited a significant increase in circRNA detection, in comparison to the matched poly-A samples, specifically a 4.6-fold increase in CIRI2 and a 22.3-fold increase in CIRCexplorer2 counts (Figure 2). The circRNA transcripts were quantified from these samples, and a dimensionality reduction was performed using UMAP. There is evident clustering by RNA-seq selection protocol, further establishing the difference in circRNA quantification between poly(A)- selected and rRNA-depleted data (Supplementary Figure 7). rRNA-depleted samples had a 1.4 – 9.6-fold increase in the normalized counts of unique circRNA transcripts detected compared to poly(A)-selected samples (Supplementary Figure 8a). When comparing transcripts detected in both poly(A)-selected and rRNA-depleted samples, there is a 0.31 – 0.86-fold increase in circRNA counts in rRNA-depleted samples (Supplementary Figure 8b-c).

**Figure 2.**
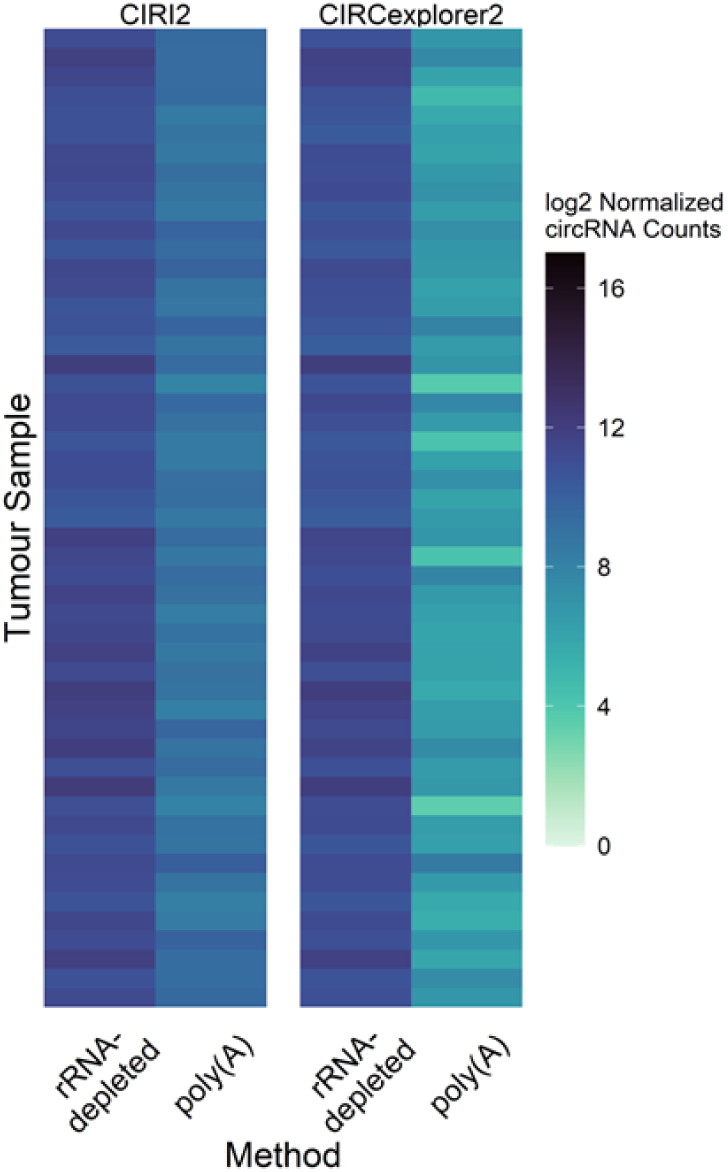
circRNA expression across 51 lung tumors with poly(A)-selection and matched rRNA-depleted samples. CIRI2 and CIRCexplorer2 used for detection. Expression is log_2_+1 normalized read counts. Only circRNAs with ≥ 2 junction reads were retained. Colour bar units represent log_2_+1 normalized count values from 0 –16, where dark blue indicates high circRNA expression, while light blue indicates low circRNA expression.

circRNA identification using rRNA-depleted samples yielded not only a higher quantification of transcripts, but also captured more unique transcripts than poly(A)-selected samples. These findings further prove that the RNA-seq selection protocol has a significant impact on the resulting circRNA profiles as previous studies have also observed (47), and brings to question the reliability of previous studies that have used the poly(A)-selection protocol on circRNA detection for biomarker discovery.

### Expression and sequencing features associated with stability of transcriptomic profiling

Identifying biological and sequencing features that may impact the stability of transcript expression will help characterize the well-documented inconsistency between pharmacogenomic studies (14, 15, 17, 18, 20, 23). To ensure the training dataset had an optimal balance between transcripts classified as either stable or unstable, the quantile of transcripts with the highest Kuncheva index was selected. The Kuncheva index was highest for the top stable transcripts at the 15% quantile, which were used to create a set of “stable” and “unstable” transcripts’ labels (Supplementary Figure 9). To identify the sequencing features that are associated with transcript stability, we trained a linear regression model to quantify the strength and significance of associations between the transcript features – median expression, length (sum of lengths of exons), number of exons, GC-content, 3’/5’ mapping bias, and gene mappability – and stability as a continuous or binary value, respectively. The linear regression model were highly significant (p-value < 2.2e-16; Table 2) and predictive (MSE < 0.133; Table 2). The importance of the transcriptomic features on the stability indices were investigated following permutations (*n*=20,000) (Supplementary Figure 10). The feature influence showed significant within features (Table 3) and across features (Supplementary Table 3). Median expression and mRNA length were the two features with the greatest influence (Figure 3).

**Figure 3.**
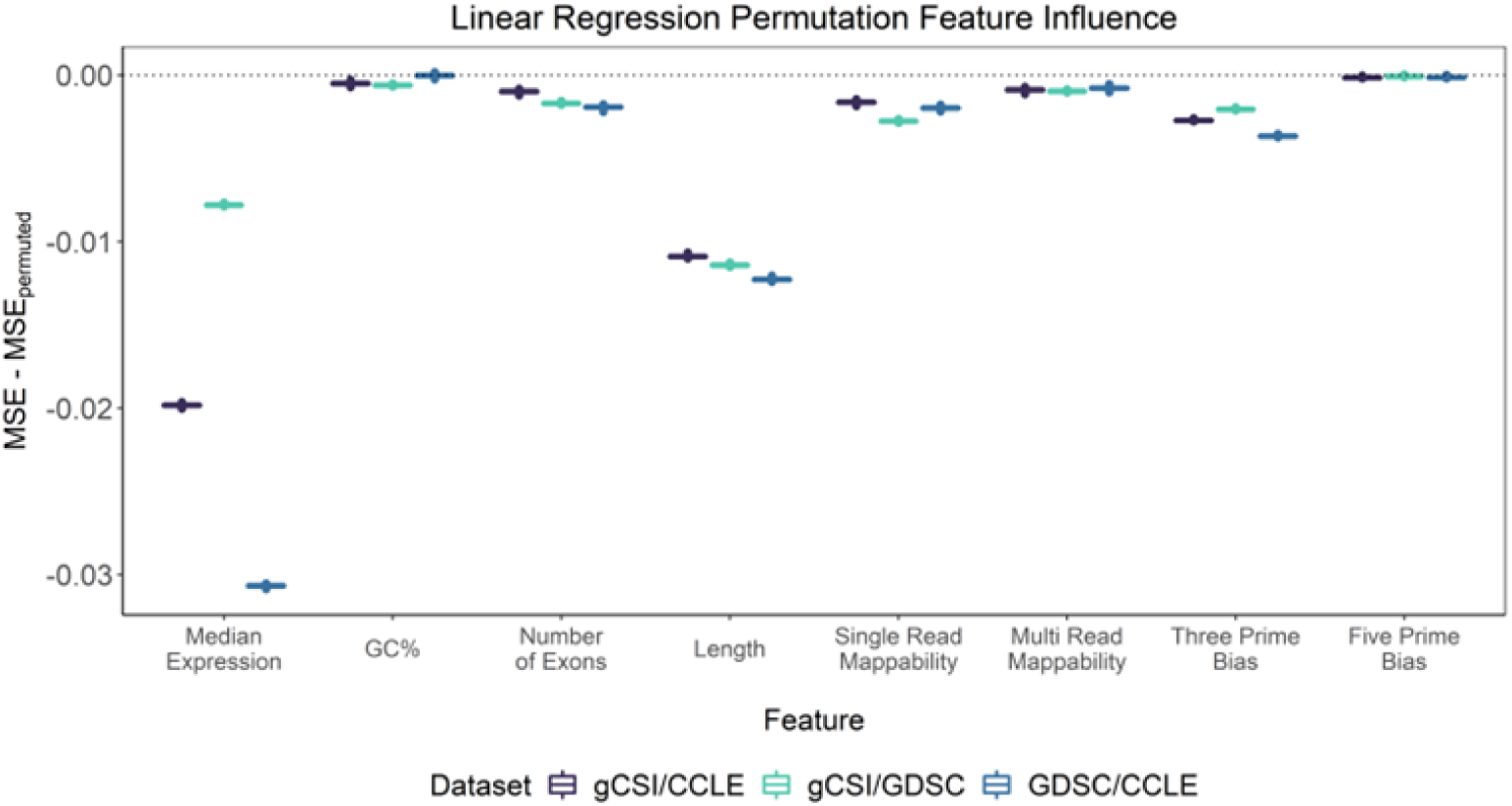
Feature influence across dataset pairs computed through comparison of baseline and permuted scoring metric (n=20,000), with 10 repetitions. Baseline (MSE) and permuted scores (MSE_permuted_) represent mean score across all folds (10-fold CV).

**Table 2.** Performance of linear regression across dataset pairs. The performance metrics are computed as the mean ± standard deviation (10-fold CV with 10 repetitions). Performance metrics not utilized for each respective algorithm are represented as NA.

| Algorithm | Dataset Pair | MSE(baseline) | Accuracy(baseline) | P-Value |
| --- | --- | --- | --- | --- |
| Linear Regression | gCSI/CCLE | 0.127±0.011 | NA | < 2.2e-16 |
|  | gCSI/GDSC2 | 0.137±0.008 | NA | < 2.2e-16 |
|  | GDSC2/CCLE | 0.136±0.014 | NA | < 2.2e-16 |

**Table 3.** Mean performance of linear regression model across dataset pairs before and after feature permutation. Metric value is the average MSE score across all folds (10-fold CV) after 10 repetitions. P-value represents the deviation of the permuted scores from the baseline.

| Algorithm | Feature | Metric(baseline) | Metric(permuted) | P-Value |
| --- | --- | --- | --- | --- |
| Linear<br>Regression | Median Expression | 0.1103694 | 0.129805 | < 2.2e-16 |
|  | GC% | 0.129806 | 0.1301876 | < 2.2e-16 |
|  | No. Exons | 0.1301888 | 0.1317269 | < 2.2e-16 |
|  | Length | 0.1347396 | 0.1462667 | < 2.2e-16 |
|  | Single Read Mappability | 0.1317286 | 0.133856 | < 2.2e-16 |
|  | Multi Read Mappability | 0.1338565 | 0.1347364 | < 2.2e-16 |
|  | 3 Prime Bias | 0.1462666 | 0.1490808 | < 2.2e-16 |
|  | 5 Prime Bias | 0.1490814 | 0.1492056 | 5.52E-05 |

### Association between mRNA stability and predictive value for drug response

RNA species, such as gene isoforms and circRNAs, have been reported to be a rich source of prognostic and predictive biomarkers (8, 9, 14). We therefore assessed whether mRNAs with stably quantified expression are more likely to be more predictive of drug response. We explored the relationship between the stability of the expression quantification of two known biomarkers and drug response, namely *ERBB2* and *EGFR* expression predictive of response to Lapatinib and Erlotinib respectively (48), and *PHB* expression shown to be associated with response to Paclitaxel in preclinical studies (49). We observed that the most stable isoforms were not consistently the most predictive for the biomarkers of interest (Figure 4). While the most stable *EGFR* transcript, known as *EGFR-201*, yielded a strong predictive value that is on par with the overall mRNA expression, the most stable transcripts for *ERBB2* and *PHB* did not follow the same pattern. By expanding the analysis to all genes, we observed that the distribution of associations between the stability of mRNA and their predictive value was not better than random for the 6 drugs in common between CCLE, GDSC2 and gCSI (Supplementary Figure 11), suggesting that mRNA stability is not a strong predictor of drug response.

**Figure 4.**
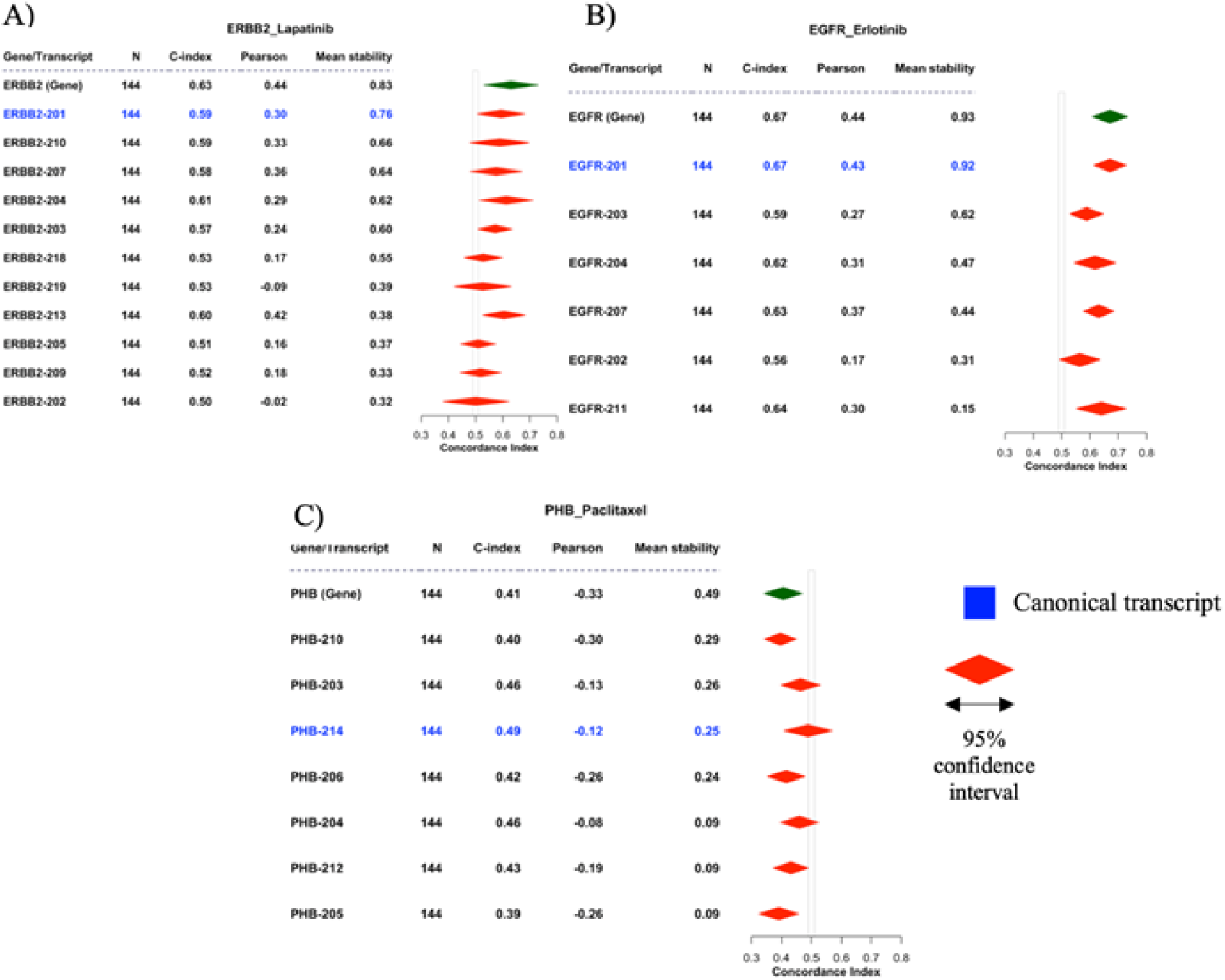
Forest plots displaying drug associations with known genes expression-based biomarkers and their respective mRNAs with corresponding stability. **A)** *ERBB2*/Lapatinib; **B)** *EGFR*/Erlotinib; **C)** *PHB*/Paclitaxel. N: sum of biological replicates across gCSI, CCLE, GDSC2; C-index: concordance-index; Pearson: Pearson correlation coefficient; mean stability: mean stability index of a mRNA across the three dataset pairs (gCSI/CCLE, gCSI/GDSC2, GDSC2/CCLE). All values computed are a meta-estimate across the dataset pairs. C-index displayed with 95% confidence interval. Canonical transcripts labeled in blue were obtained through UCSC (transcripts with the longest CDS region). GENCODE annotation used for Gene/Transcript labels. Grey bar denotes baseline CI of 0.5.

### High *hsa_circ_0001159* expression associated with prolonged patient overall survival

To investigate the prognostic value of circRNAs, we analyzed a cohort of 51 adenocarcinoma patients whose tumours have been profiled using rRNA-depletion RNA-seq. We quantified the prognostic value of each circRNA using concordance index (CI) with patients’ overall survival data (Figure 5, Supplementary Figure 12). We established a filtering criterion to retain only significant associations (CI > 0.65 or < 0.35 and FDR < 5%), in which only one circRNA, *hsa_circ_0001159*, was identified. This circRNA transcript was detected using both CIRI2 and CIRCexplorer2. The higher circRNA expression of *hsa_circ_0001159* was associated with prolonged overall survival of adenocarcinoma patients, where CIRI2 yielded a concordance-index of 0.26 (FDR = 0.034), while CIRCexplorer2 yielded a CI of 0.21 (FDR = 0.003). Additionally, *hsa_circ_0001159* expression was robust across quantification pipelines, yielding a Spearman correlation coefficient of 0.92. Therefore, these results suggest *hsa_circ_0001159* as a potential prognostic marker in lung adenocarcinoma, although its prognostic value will need to be further validated in independent datasets.

**Figure 5.**
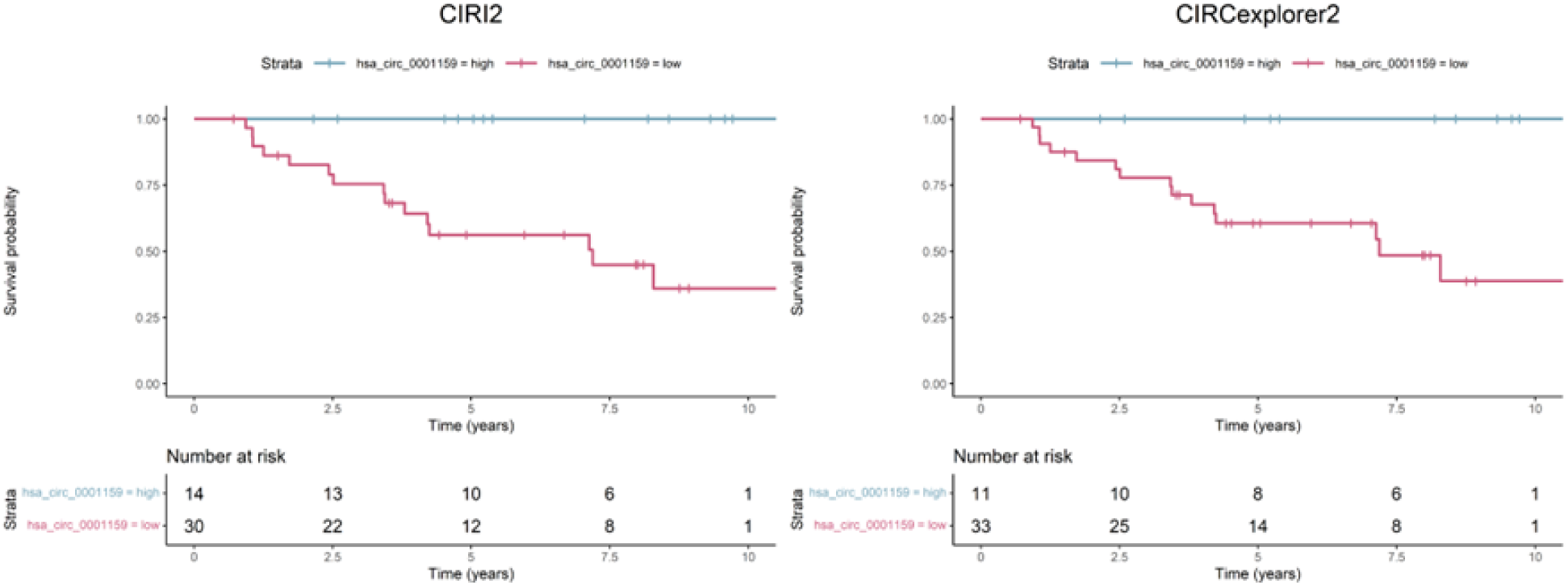
Kaplan-Meier curves displaying overall adenocarcinoma patient survival for *hsa_circ_0001159* detected by both CIRI2 and CIRCexplorer2. CI: concordance index, FDR: false-discovery-rate. rRNA-depleted profiles are used for circRNA detection. Only circRNAs with ≥ 2 junction reads are retained. Red strata depicts high expression of *hsa_circ_0001159*, while blue depicts low expression of *hsa_circ_0001159*. High expression is defined as log_2_ normalized count value of a respective circRNA ≥ 3.21, while low expression is defined as log_2_ normalized count value of a respective circRNA < 3.21. *hsa_circ_0001159* annotation derived from circBase.

## DISCUSSION

In this study, we investigated the consistency of transcript expression across biological replicates from three large pharmacogenomic studies. circRNA expression was immensely inconsistent across biological replicates, expressing that caution should be taken when using the transcriptomic data from the pharmacogenomic datasets for biomarker discovery. The lack of transcript stability suggests that a potential biomarker discovered in one dataset may not be detected in another, even though the same cell line and RNA isolation protocol is used for RNA-seq (poly(A)-selection). Moreover, investigation into mRNA biological and sequencing features led to the discovery that median expression and mRNA transcript length influence the prediction of transcript stability, which provides further insight into the consistency of mRNA expression across these datasets. The variability in signal when comparing replicates across datasets may be due to the different experiment protocols used to generate the samples (Supplementary Table 2). However, the same pattern of divergent transcript detection and expression stability observed in intra-lab comparisons aid to validate our findings. To obtain more confidence in these results, further investigation with a larger circRNA sample size is required.

Investigation into the association between mRNA stability and predictive value found no associations, including across a multitude of genes, as there was a large overlap between randomized and non-randomized correlations between mean stability and drug response of genes for intersected drugs. Therefore, the most stable mRNAs are not consistently the most predictive of drug response across the biological replicates of gCSI, CCLE, and GDSC2. As previous studies have reported the inconsistency of drug response measurements across laboratories, which may prevent the discovery of an association, if any (15, 17), further investigation may be needed before a lack of association can be concluded.

To our knowledge, the prognostic signal of *hsa_circ_0001159* that we identified in lung adenocarcinoma has not been investigated. *hsa_circ_0001159* is expressed by the Chromodomain Helicase DNA Binding Protein 6 (CHD6) gene (44), where high expression of CHD6 has been associated with greater survival in adenocarcinoma patients by the Human Protein Atlas (www.proteinatlas.org). Moreover, CHD6 has also been identified as regulating oxidative DNA damage response (50). However, the biological function and potential association of *hsa_circ_0001159* with adenocarcinoma needs to be further investigated through circRNA validation with RNAse-R/RT-PCR, including in additional patient cohorts.

We utilized RNA-seq data with both RiboZero and RiboMinus rRNA-depletion protocols, where RiboZero and RiboMinus removes 5S, 5.8S, 18S, and 28S rRNA from a sample, however RiboMinus is a product of Thermo Fisher Scientific, while RiboZero is a product of Illumina. In addition, RiboZero has been shown to increase rRNA removal, when compared to a RiboMinus (34), suggesting a potential influence of depletion kits on non-coding detection. Therefore, further investigation is needed into the impact of various depletion kits on circRNA detection. We used Kallisto for mRNA and gene quantification, a tool that pseudo-aligns to a reference transcriptome for rapid expression quantification (51–53). However, this can be limiting, as the tool is unable to detect novel isoforms outside of a provided transcriptome, which impacts the biomarkers discovered. CIRI2 is a known circRNA detection tool that utilizes efficient maximum likelihood estimation (MLE) for back-splice junction identification from RNA-seq data (54), while CIRCexplorer2, another known tool for circRNA detection, utilizes unmapped reads from TopHat as input into TopHat-Fusion to identify back-splice junction reads on the same chromosome, which also allows for de novo assembly and alternative splicing event detections (55, 56). CIRI2 and CIRCexplorer2 performance have both been benchmarked, where CIRI2 has been shown to have a greater sensitivity and precision than CIRCexplorer2 for detection in a dataset with only circRNA species, while CIRCexplorer2 was found to have greater precision with mixed datasets (57). Therefore, both of these tools were used in our study for circRNA detection, in order to address any software bias, as a potential biomarker may be discovered using one tool but not the other. Overall, we invalidated the use of poly(A)-selected RNA-seq data from pharmacogenomic consortia for circRNA detection and drug response associations from other studies (7, 12), through biological replicates and identification of a significant detection decrease when validated with matched RNAse R treated samples. This also displays the significance of incorporating matched RNAse R samples in order to validate circRNAs detected, even if a rRNA-depleted method such as RiboMinus is utilized.

## DATA AVAILABILITY

All pharmacogenomic datasets and their respective RNA-seq and drug sensitivity data utilized in the analyses can be retrieved as data objects from the ORCESTRA platform (www.orcestra.ca). The data objects were analyzed in R (v.4.2.1) using the *PharmacoGx* (v.3.0.2) Bioconductor package. The Zenodo DOI’s for each object are: gCSI (10.5281/zenodo.4737437); CCLE (10.5281/zenodo.3905461); GDSC2 (10.5281/zenodo.5787145). The data for each analysis can be accessed via our custom compute capsule on Code Ocean (58) (<u>codeocean.com/capsule/1660304/tree</u>).

## CODE AVAILABILITY

To ensure full reproducibility and transparency of our results, we utilized Snakemake (<u>snakemake.readthedocs.io/</u>), a workflow-management tool that allows for the reproducible execution of our CIRI2 and CIRCexplorer2 pipelines for circRNA detection. They can be accessed on Code Ocean (58), where our code can be executed through a custom compute environment, ensuring reproducibility and transparency for our findings (<u>codeocean.com/capsule/1660304/tree</u>). All code is available on GitHub (https://github.com/bhklab/circRNA-detection).

## ACKNOWLEDGEMENTS

This project is supported by the Canadian Institutes of Health Research (CIHR), under the frame of ERA PerMed.

## SUPPLEMENTARY FIGURES

**Supplementary Figure 1.**
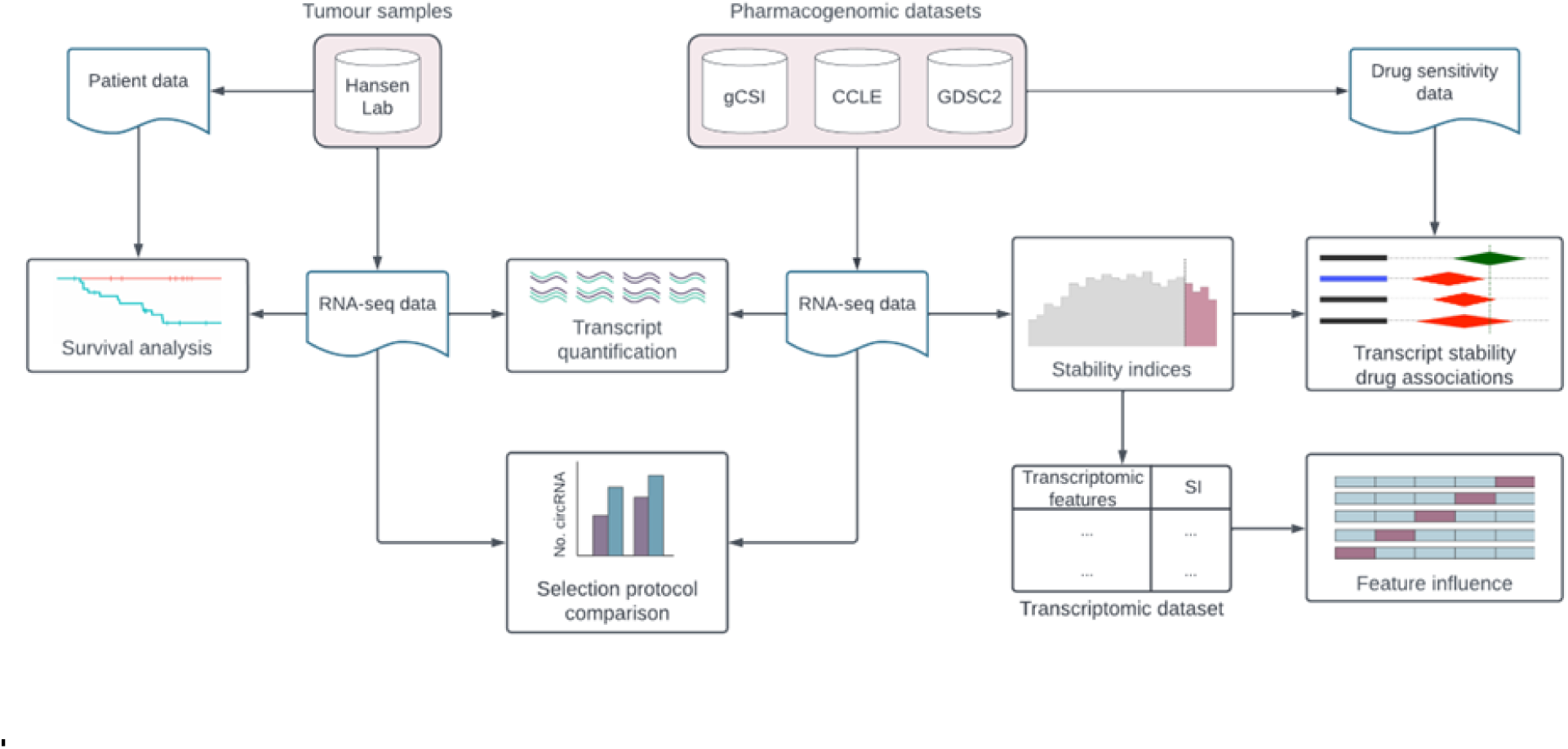
Schematic of workflow used. Datasets and samples are coloured in pink boxes.

**Supplementary Figure 2.**
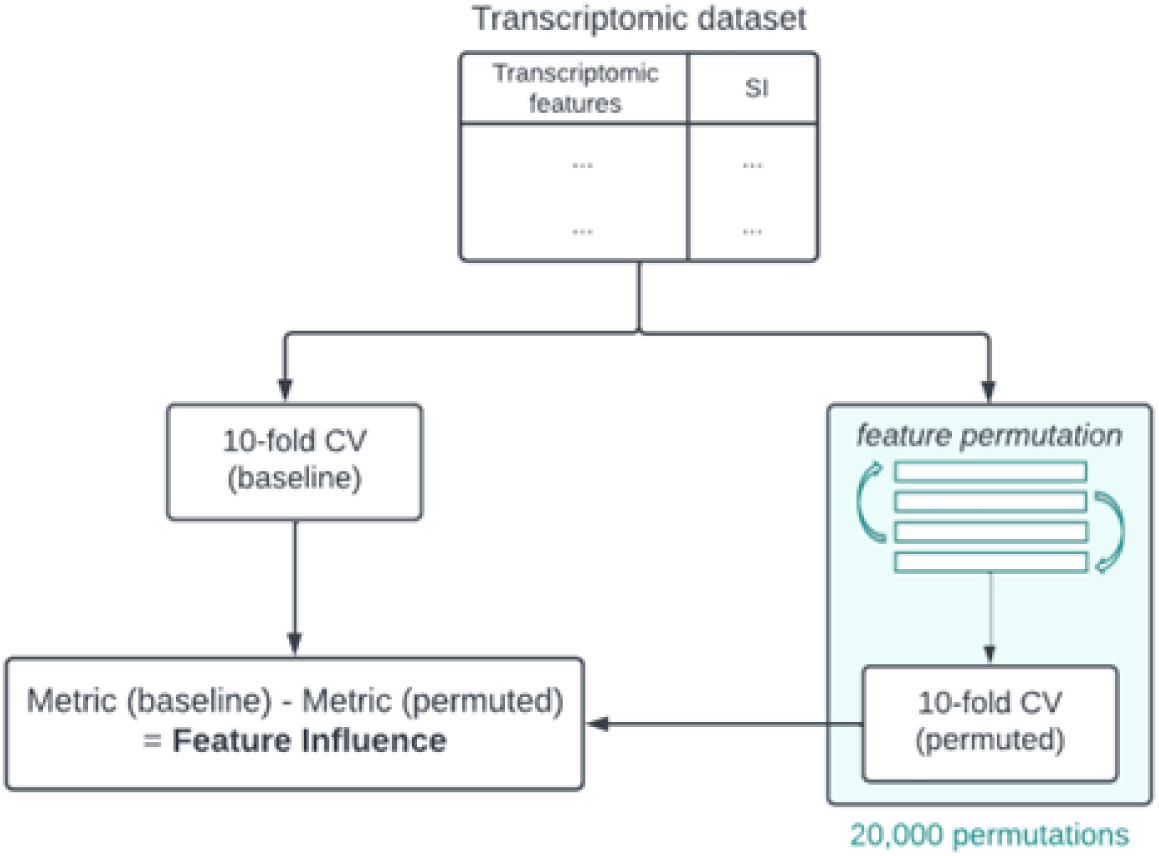
Schematic of permutation feature influence. 10-fold CV performed on training data set before and after feature permutation (n=20,000). Metric value is the average MSE score across all folds (10-fold CV) after 10 repetitions.

**Supplementary Figure 3.**
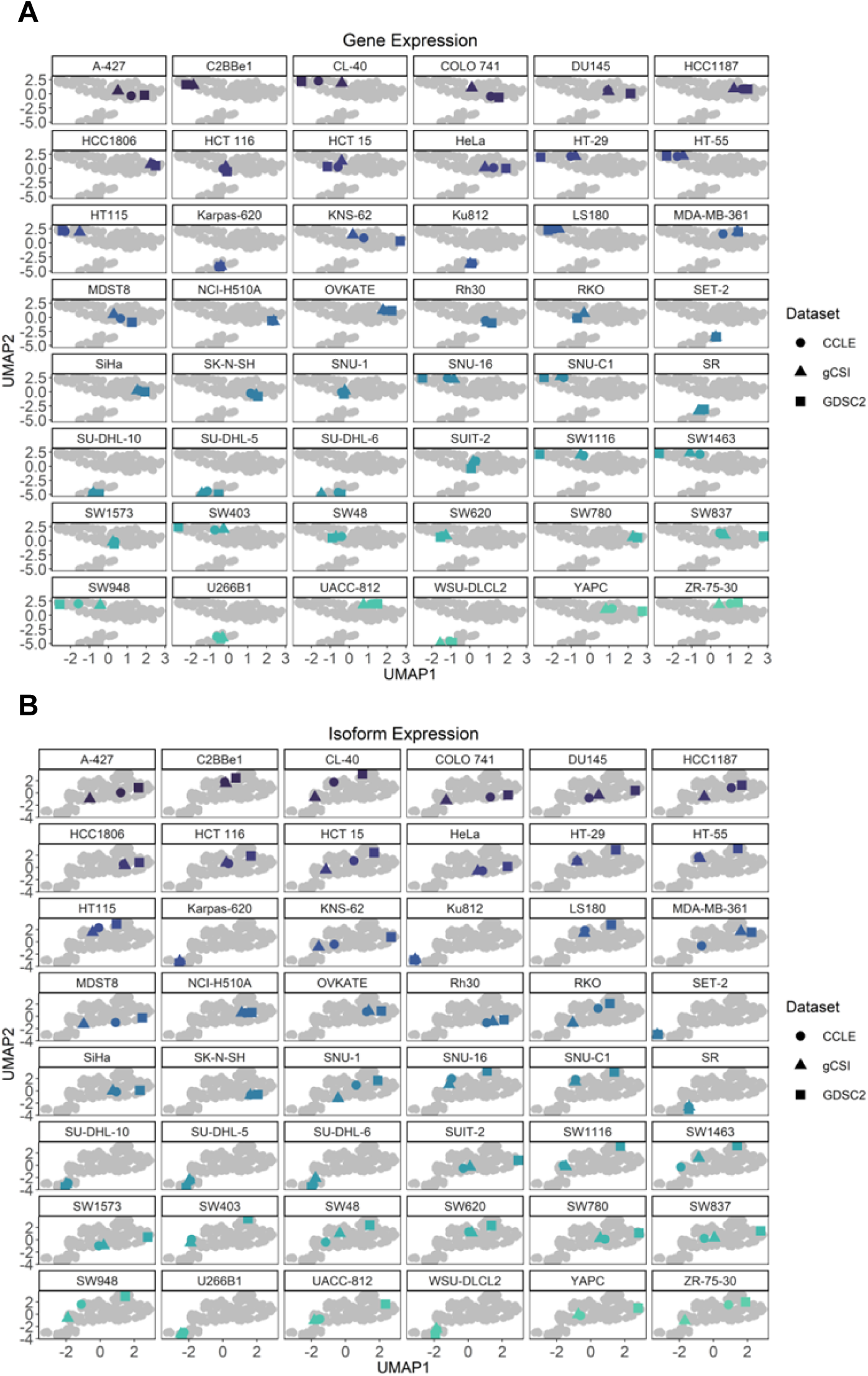
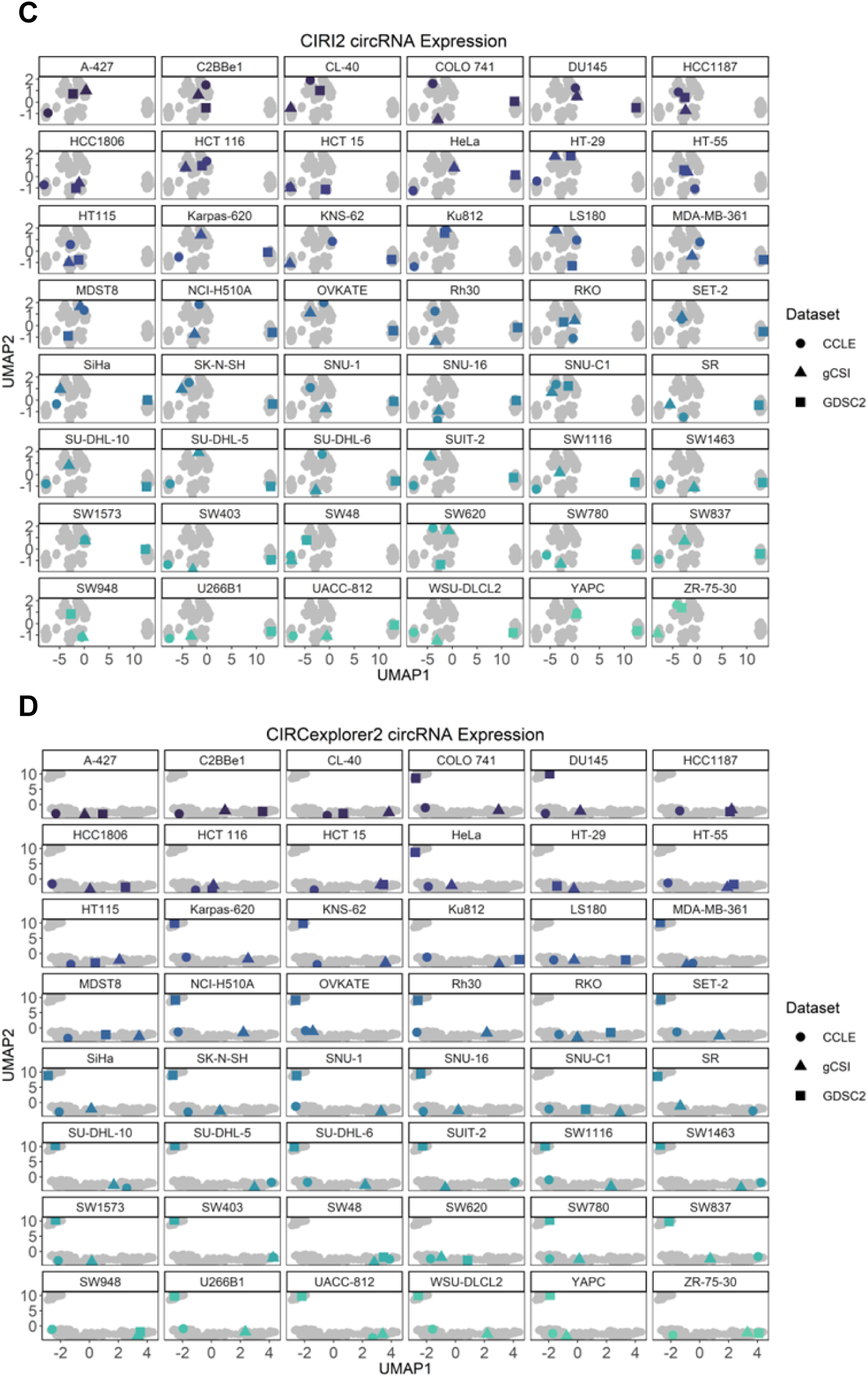
Transcript expression data dimensionality reduction by UMAP grouped into facets for each of the 48 biological replicates from 3 datasets (gCSI, CCLE, GDSC2). Expression is log_2_+1 normalized read counts. Coloured data points correspond to the specified cell line of the facet. Shape corresponds to each dataset. **A)** Gene expression data. **B)** Isoform expression data. **C)** circRNA expression data quantified using CIRI2. **D)** circRNA expression data quantified using CIRCexplorer2.

**Supplementary Figure 4.**
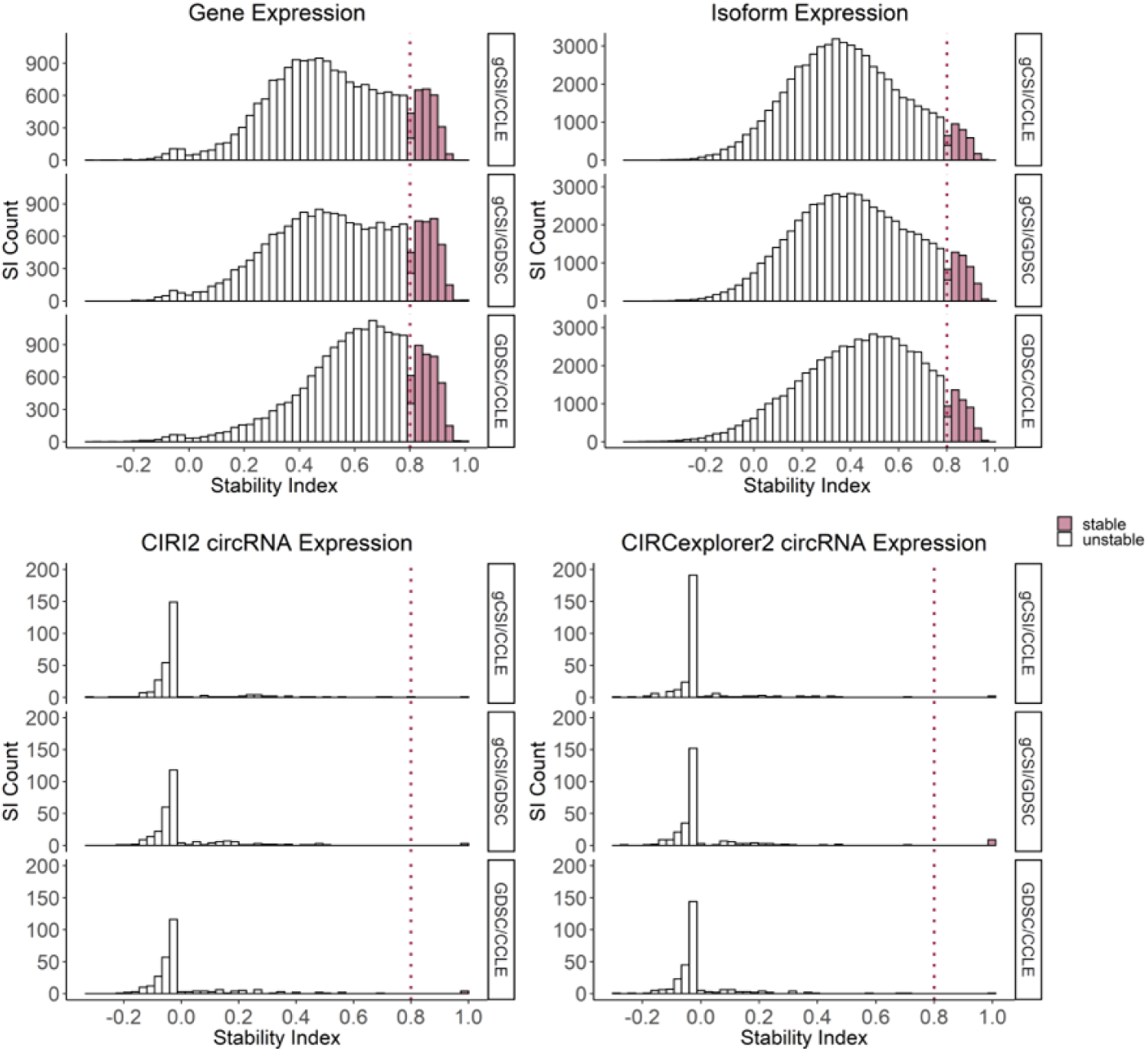
Distribution of circRNA, mRNA and corresponding gene expression stability across dataset pairs. Stability index estimated using the Spearman correlation coefficient. Coloured bars represent transcripts classified as stable (SI > 0.8).

**Supplementary Figure 5.**
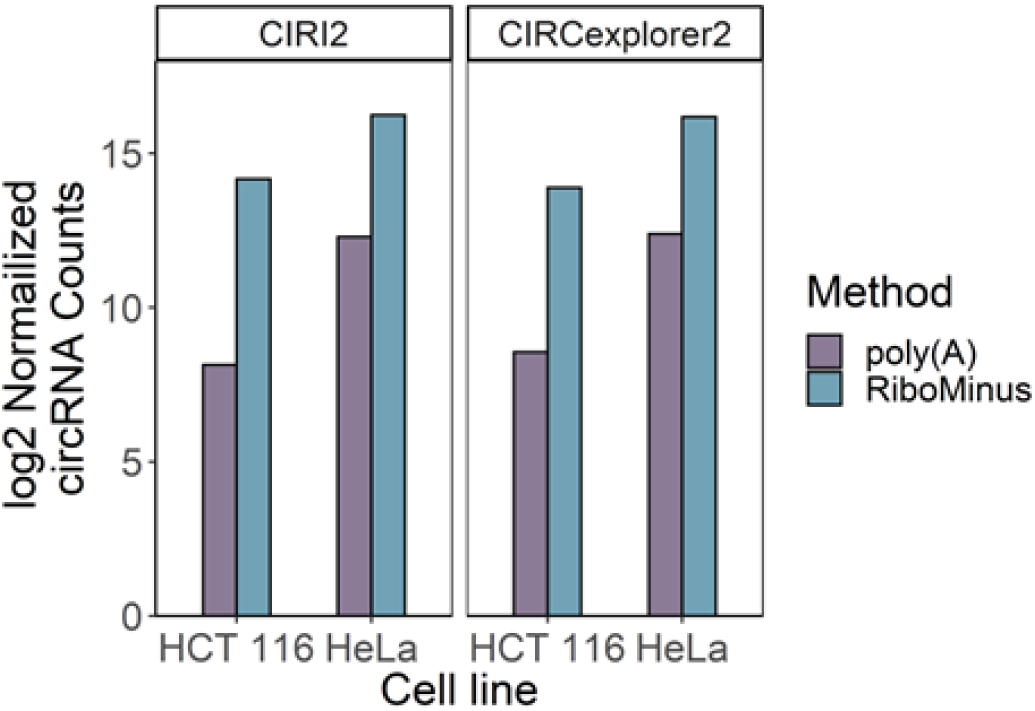
Number of circRNAs detected in each cell line with poly(A) and RiboZero selection. CIRI2 and CIRCexplorer2 used for detection. Only circRNAs with ≥ 2 junction reads are retained. Expression is log_2_ normalized read counts.

**Supplementary Figure 6.**
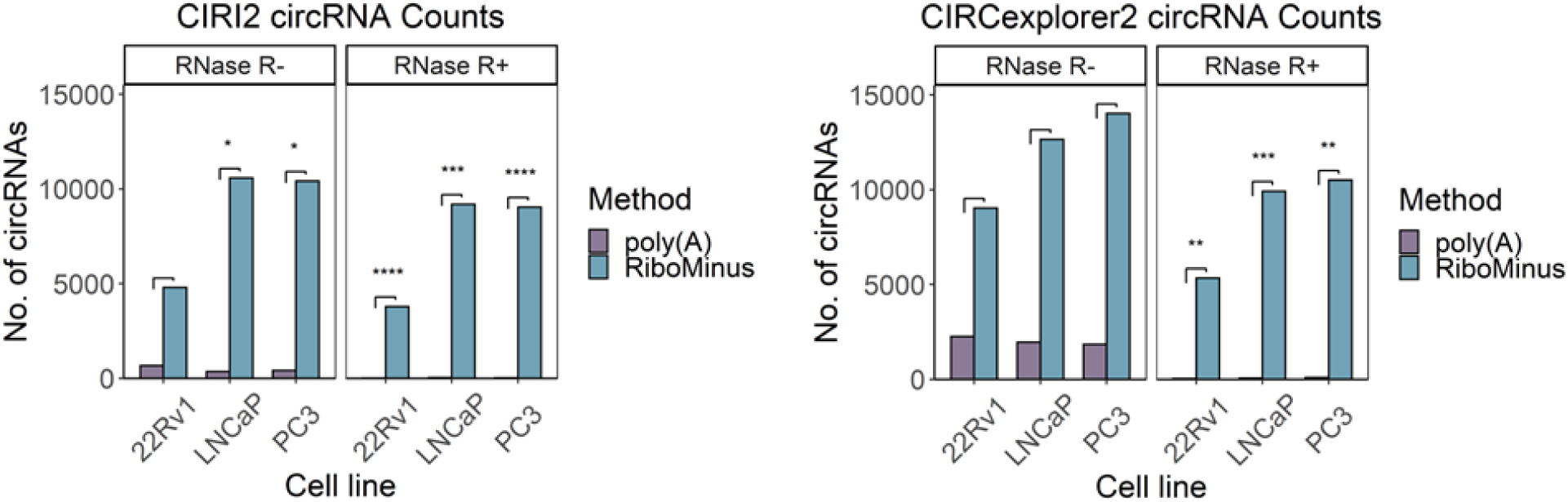
Number of circRNAs detected in each cell line with poly(A) and RiboMinus selection before and after RNAse R validation. RNAse R-: without RNAse R treatment; RNAse R+: with RNAse R treatment. Only circRNAs with ≥ 2 junction reads and at least 1.5-fold enriched in matched RNAse R+ samples are retained. No. of circRNAs are normalized via Counts per Million (CPM). * >10-fold change, ** >50-fold change, *** >150-fold change, **** >300-fold change.

**Supplementary Figure 7.**
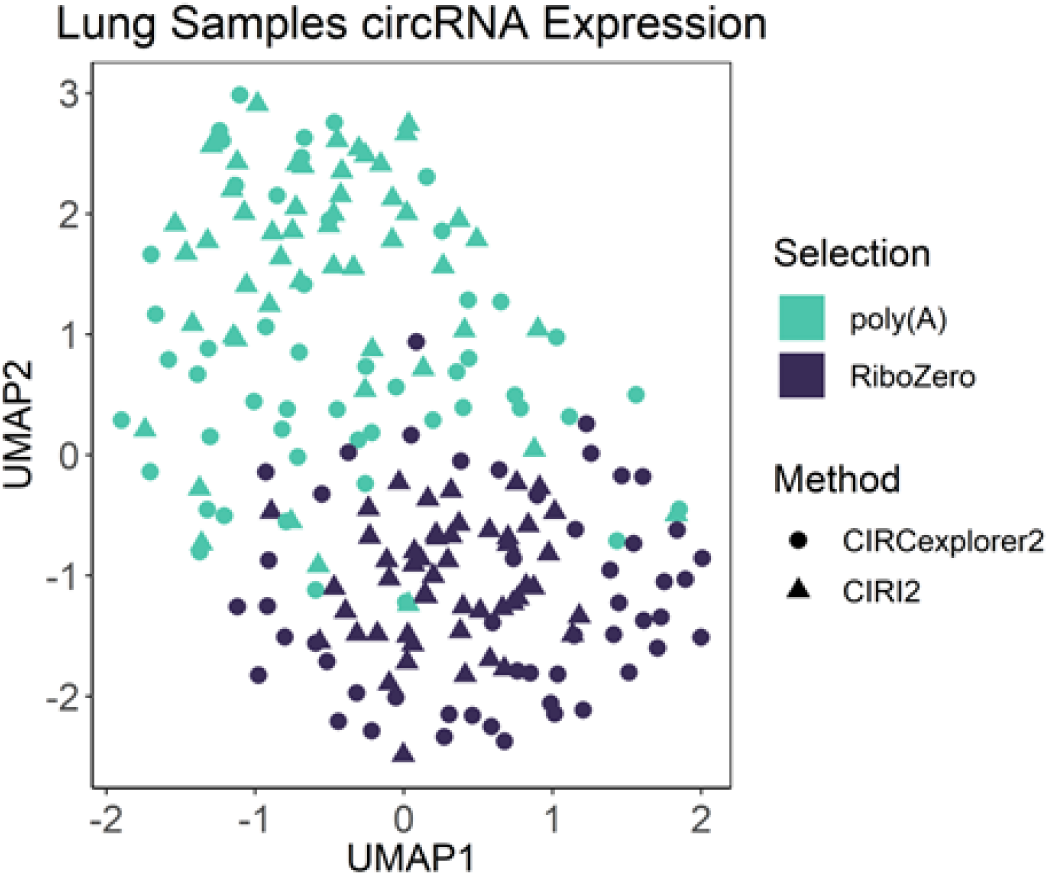
circRNA transcript expression data dimensionality reduction by UMAP. circRNA transcript expression across 51 adenocarcinoma tumour samples. Expression is log_2_+1 normalized read counts. Colour by RNA-seq selection protocol, shape by circRNA detection tool.

**Supplementary Figure 8.**
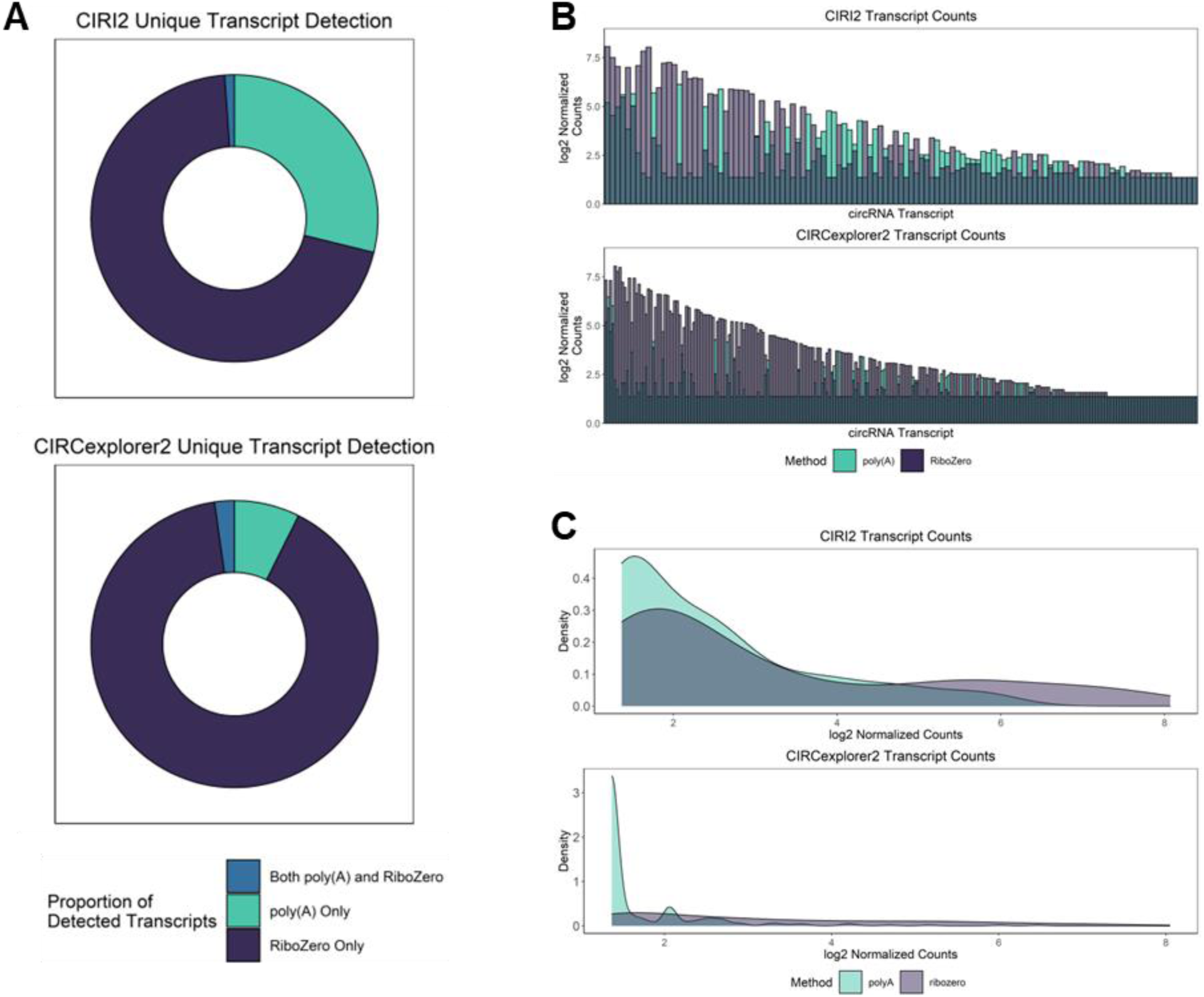
A) Proportion of unique circRNA transcripts detected across 51 lung tumors from poly(A)-selected samples, rRNA-depleted samples, or both. CIRI2 and CIRCexplorer2 used for detection. **B)** Quantification of circRNA transcripts found in both poly(A)- selection and rRNA-depleted lung tumors samples. CIRI2 and CIRCexplorer2 used for detection. Expression is log_2_+1 normalized read counts. **C)** Distribution of log_2_+1 normalized transcript counts found in both poly(A)-selection and rRNA-depleted lung tumors samples. CIRI2 and CIRCexplorer2 used for detection.

**Supplementary Figure 9.**
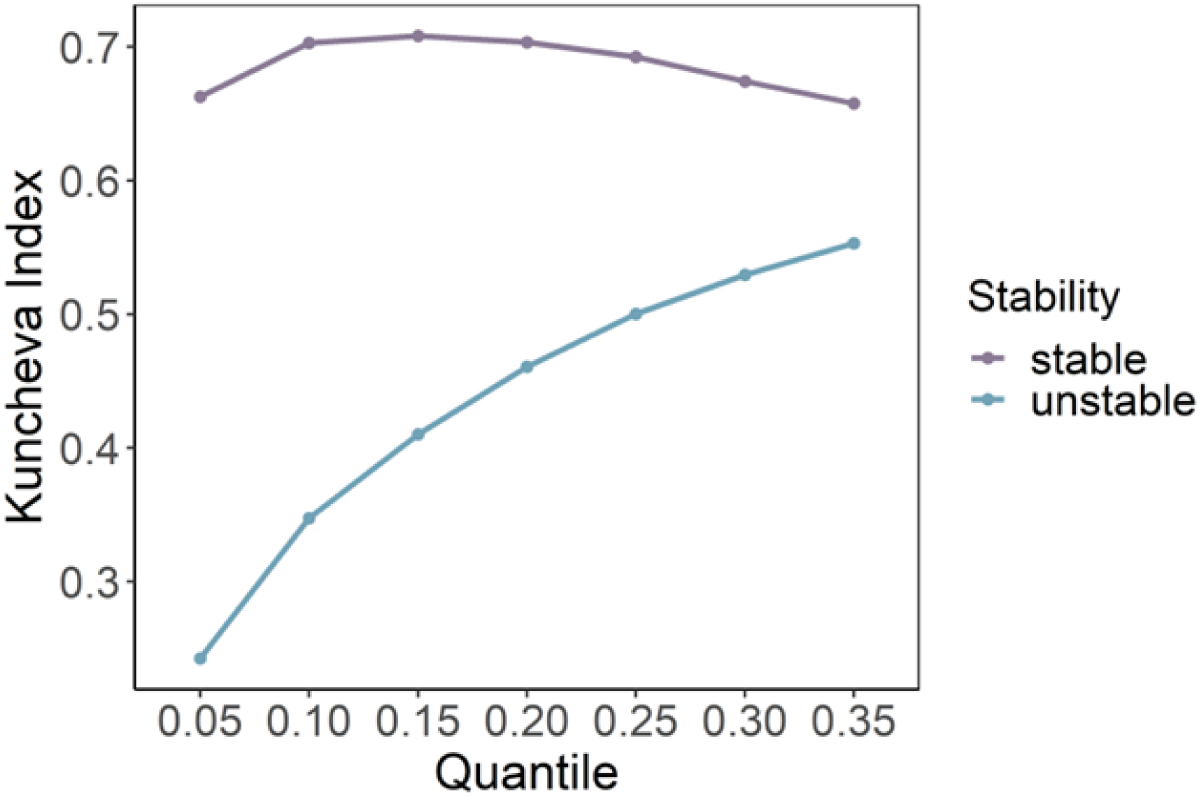
Kuncheva index computed for top stable and unstable mRNA across quantile range. Stability computed using the stability index. mRNA transcripts that possess a stability index > 0.8 classified as stable, otherwise transcript classified as unstable.

**Supplementary Figure 10.**
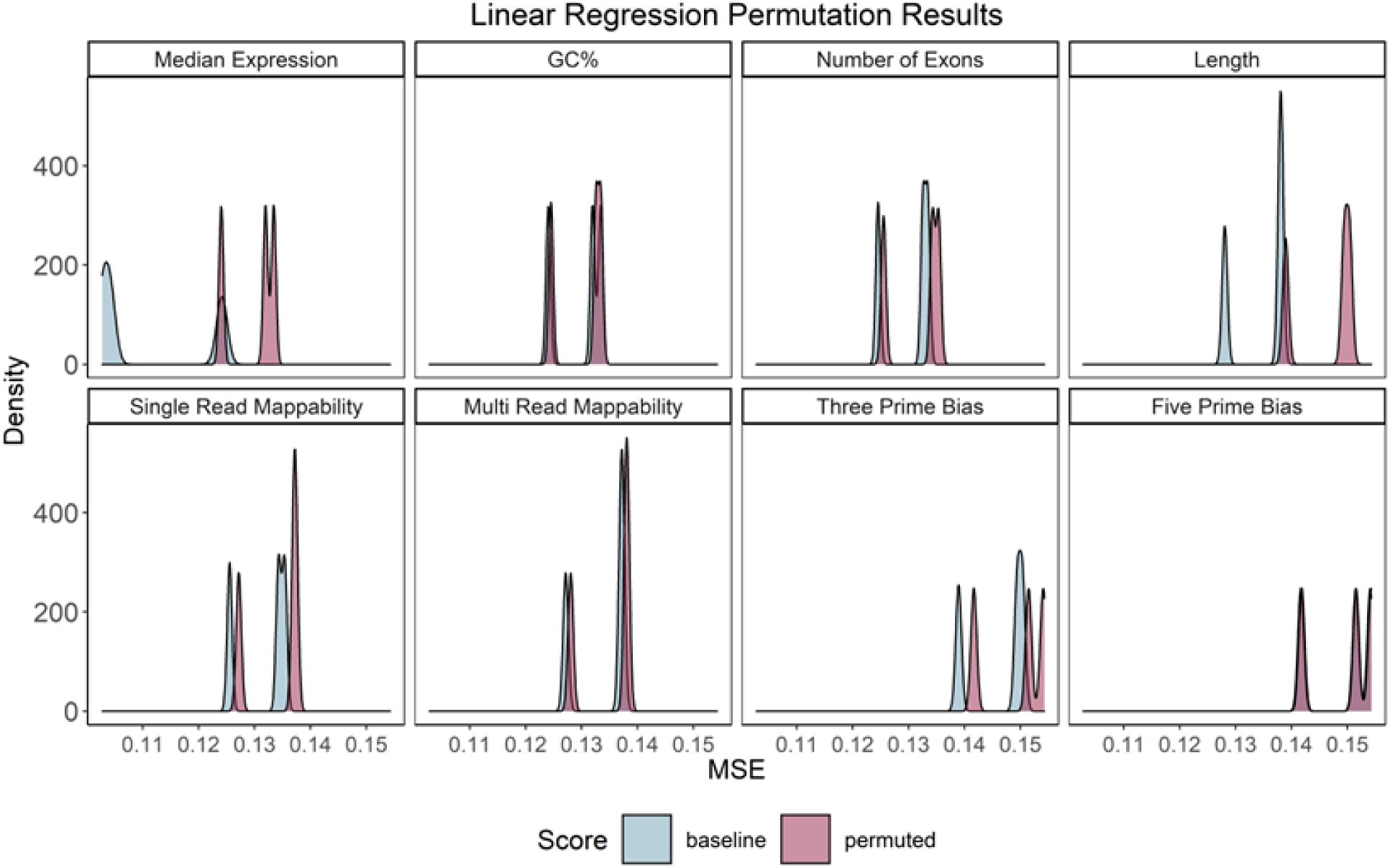
Computation of performance metric before and after feature permutation of each transcriptomic feature (n=20,000) per dataset pair. Metric value is the average MSE score across all folds (10-fold CV) after 10 repetitions.

**Supplementary Figure 11.**
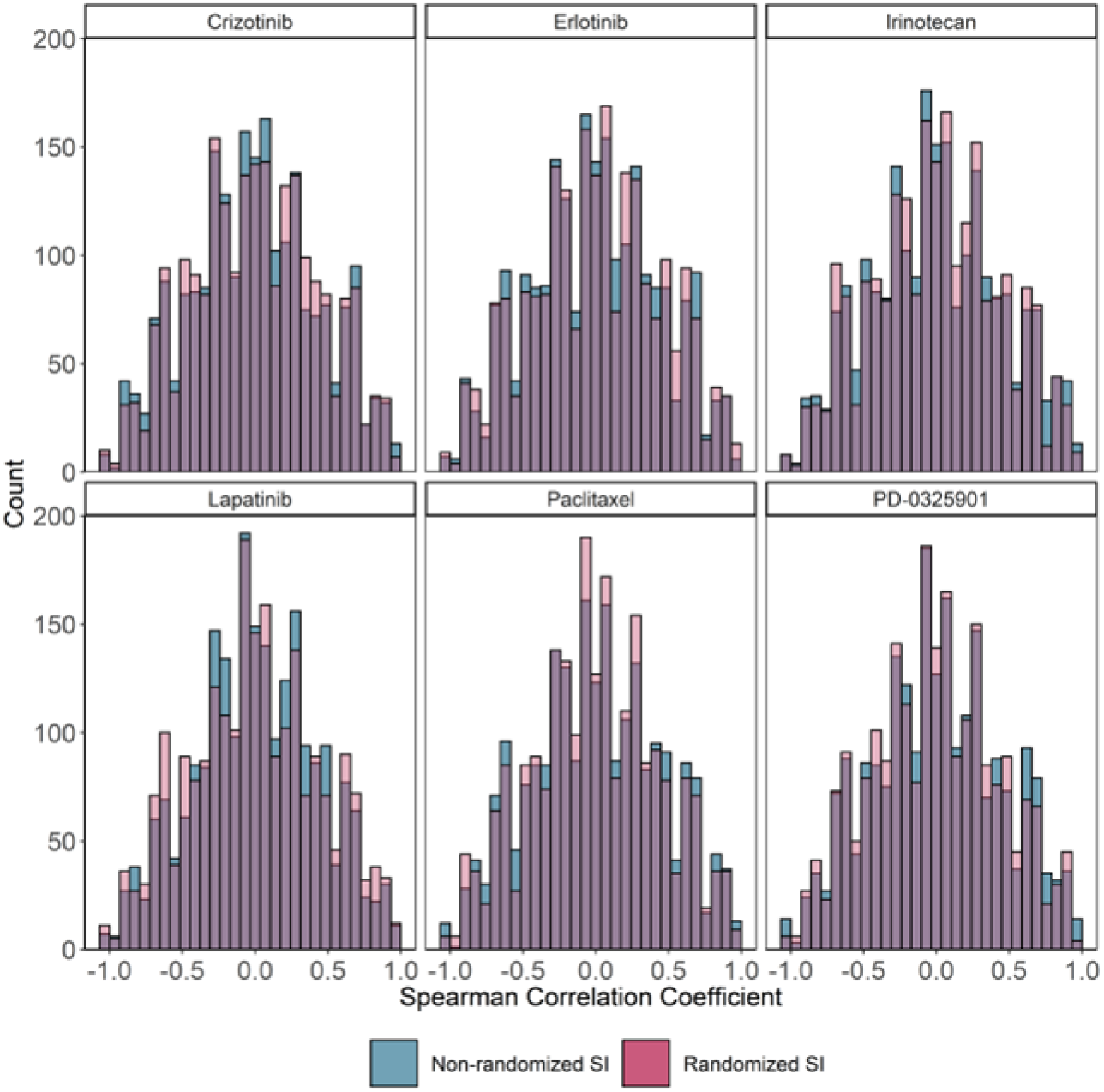
Histograms displaying Spearman correlation between mean stability and Pearson correlation coefficient across mRNAs for each gene and respective drug. SI: stability index. Spearman correlation coefficient with a randomized and non-randomized mean SI is reported. Only intersected drugs across gCSI, CCLE, and GDSC2 are selected for.

**Supplementary Figure 12.**
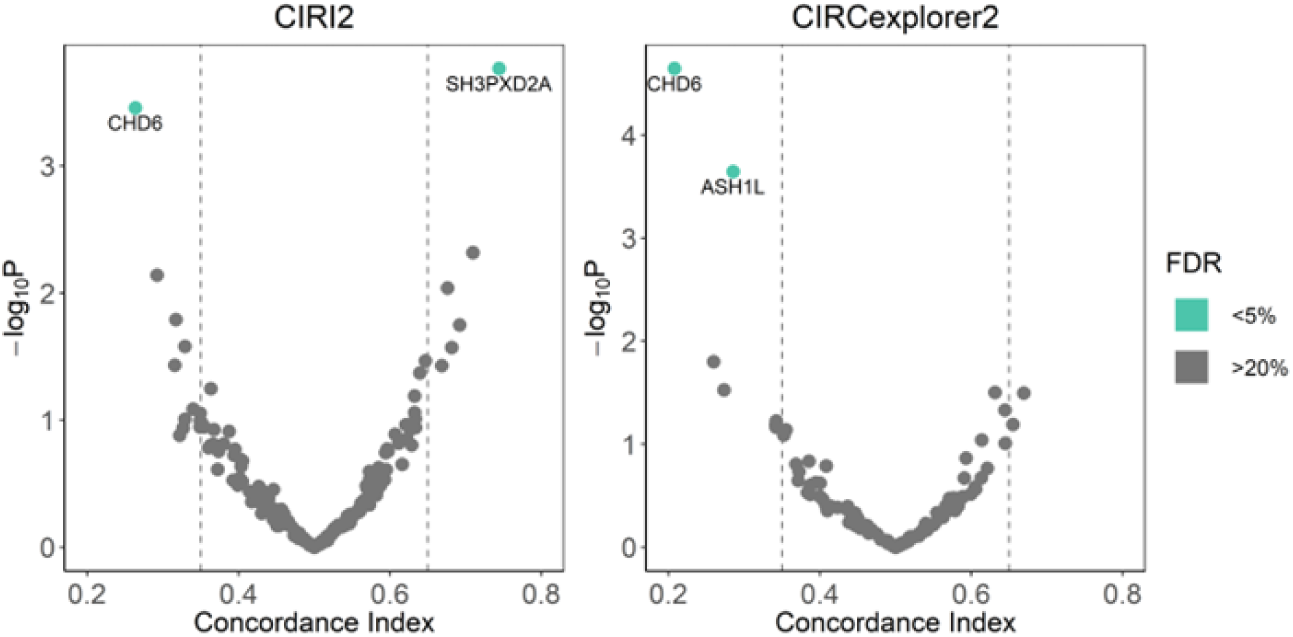
Volcano plots displaying overall adenocarcinoma patient survival for circRNAs detected by both CIRI2 and CIRCexplorer2. -log10P: -log10p-value, FDR: false-discovery-rate. Dashed vertical lines indicate concordance index thresholds (0.35 and 0.65). Green colour represents circRNAs with FDR < 5%. rRNA-depleted profiles are used for circRNA detection. Only circRNAs with ≥ 2 junction reads are retained. Selected circRNA (CI > 0.65 or < 0.35 and FDR < 5%) labeled with gene annotation by circBase.

## SUPPLEMENTARY TABLES

**Supplementary Table 1.** ORCESTRA pharmacogenomic data objects with respective cell lines, drugs, tissues, sensitivity experiments, and molecular profiles. Sensitivity experiments possess varying pharmacological assays. The Zenodo DOI’s for each object are: gCSI (10.5281/zenodo.3905452); CCLE (10.5281/zenodo.3905462); GDSC2 (10.5281/zenodo.3905481).

| Dataset | No. of cell lines | No. of drugs | No. of tissues | No. of drug sensitivity experiments | Pharmacological assay | Molecular profiles available |
| --- | --- | --- | --- | --- | --- | --- |
| gCSI | 747 | 16 | 27 | 6471 | CellTiter-Glo | Expression (RNA-seq), Mutation, CNV |
| CCLE | 1094 | 24 | 25 | 11670 | CellTiter-Glo | Expression (RNA-seq/Microarray), Mutation, CNV, Fusion |
| GDSC2 | 1104 | 190 | 29 | 215780 | CellTiter-Glo | Expression (RNA-seq/Microarray), Mutation, CNV, Fusion |

**Supplementary Table 2.** Summary of RNA-seq datasets for circRNA detection with varying library preparation and RNA selection protocols. The respective publications for each dataset are: 1) gCSI; 2) CCLE; 3) GDSC2; 4) 22Rv1, LNCaP, PC3; 5) HCT116; 6) HeLa; 7) & 8) private datasets not currently published.

| Dataset | No. of samples | RNA-seq instrument | Library preparation | RNA selection protocol | Library layout | Read length (bp) | No. of reads (million) |
| --- | --- | --- | --- | --- | --- | --- | --- |
| gCSI | 48 | Illumina HiSeq 2000 | Illumina TruSeq RNA Sample Preparation Kit | Poly-A | Paired-end | 75 | ~60 |
| CCLE | 48 | Illumina HiSeq 2000 | Illumina TruSeq RNA Sample Preparation Kit - automated variant | Poly-A |  | 101 | ~80 |
| GDSC2 | 48 | Illumina HiSeq 2000 | Strand-specific (dUTP) | Poly-A |  | 75 | ~100 |
| 22Rv1, LNCaP, PC3 (CCLE) | 3 | Illumina HiSeq 2000 | Illumina TruSeq RNA Sample Preparation Kit - automated variant | Poly-A |  | 101 | ~80 |
| 22Rv1, LNCaP, PC3 (GEO: GSE113124) | 3 | Illumina HiSeq 2000 | Illumina TruSeq Stranded mRNA Library Prep Kit with Illumina RiboMinus Human/Mouse Transcriptome Isolation kit | RiboMinus/RNase-R (+/-) |  | 125-126 | ~50 |
| HCT116 (SRR2969249) | 1 | Illumina HiSeq 2000 | Illumina TruSeq Stranded Total RNA with Illumina Ribo-Zero Human/Mouse/Rat Sample Preparation kit | RiboZero |  | 100 | ~100 |
| HeLa (SRR5413180) | 1 | Illumina HiSeq X Ten | Illumina TruSeq Stranded Total RNA HT/LT Sample Prep Kit with Epicentre Ribo-Zero rRNA Removal Kit | RiboZero |  | 150 | ~60 |
| Lung Patient Tumor | 51 | Illumina HiSeq 2500 | Illumina TruSeq RNA Library Prep Kit v2 | Poly-A |  | 125 | ~60 |
| Lung Patient Tumor | 51 | Illumina NextSeq 500 | Illumina TruSeq RNA Sample Prep Kit v2 with SMARTer Stranded Total RNA-Seq Kit v2 - Pico Input Mammalian | Ribo-depletion (SMARTer kit) |  | 76 | ~40 |

**Supplementary Table 3.** Pairwise Wilcoxon Rank Sum Tests for feature influence across features for the linear regression model corrected for multiple testing. Benjamini & Hochberg adjustment methods used. Gray boxes indicate repeated comparisons.

|  | Five Prime Bias | GC% | Length | Median | Multi Read Mappability | Number of Exons | Single Read Mappability |
| --- | --- | --- | --- | --- | --- | --- | --- |
| GC% | <2e-16 |  |  |  |  |  |  |
| Length | <2e-16 | <2e-16 |  |  |  |  |  |
| Median | <2e-16 | <2e-16 | <2e-16 |  |  |  |  |
| Multi Read Mappability | <2e-16 | <2e-16 | <2e-16 | <2e-16 |  |  |  |
| Number of Exons | <2e-16 | <2e-16 | <2e-16 | <2e-16 | <2e-16 |  |  |
| Single Read Mappability | <2e-16 | <2e-16 | <2e-16 | <2e-16 | <2e-16 | <2e-16 |  |
| Three Prime Bias | <2e-16 | <2e-16 | <2e-16 | <2e-16 | <2e-16 | <2e-16 | <2e-16 |

